# The model alga *Chlamydomonas reinhardtii* forms mutualistic interactions with *Verticillium* fungi

**DOI:** 10.64898/2026.05.20.726472

**Authors:** Zoe Prockl, Urs Neubacher, Hanna Rovenich

## Abstract

Algal–fungal interactions are widespread in natural environments and contribute to key ecological processes such as primary production, nutrient cycling, and soil formation. Yet, molecular mechanisms underlying these associations remain poorly understood due to the complexity of natural systems and the lack of experimentally tractable models. Here, we describe a genetically tractable interaction between the model alga *Chlamydomonas reinhardtii* and the soil-borne fungus *Verticillium dahliae*. We show that these organisms form a contact-associated mutualistic interaction characterized by increased algal and fungal biomass. The algal physiological benefits persist under abiotic and microbial stress conditions. Across multiple *Verticillium* species, the interaction consistently promotes *C. reinhardtii* growth. In contrast, interaction outcomes vary across different *Chlamydomonas* species, where mutualistic interactions appear restricted to *C. reinhardtii*. Together, these findings demonstrate that algal–fungal interactions are shaped by asymmetric partner specificity and establish a tractable experimental system to dissect the mechanisms underlying partner compatibility in algal–fungal associations.

## INTRODUCTION

Organisms in natural environments rarely occur in isolation; instead, they engage in diverse interactions that shape ecological communities and evolutionary trajectories (1,2). Such interactions can have beneficial, neutral, or detrimental effects on the partners involved and are widespread among microorganisms inhabiting terrestrial ecosystems.

Interactions between fungi and photosynthetic microorganisms occur in a variety of natural environments (3). The most well-known examples are lichens, in which a fungal partner forms a stable mutualistic association with a photosynthetic photobiont, typically a chlorophyte alga and/or a cyanobacterium (4). These interactions are characterized by close physical contact between both partners and the formation of a stratified three-dimensional hyphal structure in which photobiont cells are embedded. Within this microenvironment, photobiont cells are protected from environmental stresses common in terrestrial habitats, including UV irradiation and drought (5). However, algal–fungal interactions are not restricted to lichen symbioses. Other algal–fungal associations have also been described (6). For example, a recent study reported an additional form of association termed alcobiosis, in which crusts formed by wood- or bark-dwelling basidiomycete fungi contain a layer of photosynthetically active algal cells (7). Moreover, environmental surveys increasingly reveal frequent co-occurrence of algae and fungi in terrestrial habitats such as soils, biological soil crusts, rock surfaces, and plant-associated microbiomes (8–12), suggesting that algal–fungal interactions may be more widespread than currently recognized.

Despite the apparent prevalence of algal–fungal associations in nature, the cellular and molecular mechanisms that govern these interactions remain poorly understood. Natural systems are often complex and involve multiple microbial partners, which makes it difficult to isolate and experimentally manipulate the interaction between specific organisms (13). In addition, many of the organisms involved in these associations cannot be readily cultivated axenically under laboratory conditions or lack genetic tools that would enable mechanistic investigation. Thus, mechanistic research on interactions between green algae and fungi is rare (14). These limitations highlight the need for experimentally tractable model systems in which algal–fungal interactions can be studied under controlled conditions, facilitating the investigation of underlying molecular mechanisms.

In recent years, experimental algal–fungal systems have revealed that interactions between algae and fungi can result in mutualistic outcomes under defined laboratory conditions, often mediated by nutrient exchange (15–17). For example, in association with *Saccharomyces cerevisiae*, growth of *Chlamydomonas reinhardtii* is supported by fungal respiration, which increases CO_2_ availability, while the alga provides reduced nitrogen in the form of ammonia (15). Similar interactions have been observed with filamentous fungi, and in some cases lead to particularly intimate associations, such as the incorporation of metabolically active cells of the marine alga *Nannochloropsis oceanica* into hyphae of the soil-dwelling fungus *Mortierella elongata* (18). In addition to nutrient exchange, protective effects have been described, as exemplified by an artificial lichen-like association in which *Aspergillus nidulans* protects *C. reinhardtii* from the effects of a bacterial toxic lipopeptide (19). Together, these studies demonstrate that algae and fungi can form stable and functionally diverse associations under laboratory conditions, highlighting the potential of synthetic systems to explore interkingdom interactions. However, most existing systems have primarily been developed to study metabolic cooperation or for biotechnological applications and have not been widely adopted as genetically tractable models for investigating the ecological and mechanistic basis of algal–fungal interactions.

Here, we establish a genetically tractable algal–fungal model system based on the alga *C. reinhardtii*, which dwells in freshwater and soil habitats, and the soil-borne filamentous fungus *Verticillium dahliae*. As a well-established experimental organism with extensive genetic and molecular tools, *C. reinhardtii* is particularly well suited for investigating the cellular and physiological responses of algal hosts during microbial interactions (20). *V. dahliae* is best known as a vascular plant pathogen with a broad host range but also interacts with diverse microbiota in both host-associated and soil environments (21). Using these organisms, we investigate the establishment, functional consequences, and variability of algal–fungal interactions under controlled conditions, with the aim of identifying the factors that govern interaction outcomes between these partners.

## MATERIALS AND METHODS

### Organisms and culture conditions

The fungal strains used in this study comprised *Verticillium dahliae* JR2, VdLs17, CQ2, TO22, 85S, V700, DVD3, and Vd39. In addition, other *Verticillium* species were included: *V. nonalfalfae* T2, *V. alfalfae* PD638, *V. nubilum* 397, *V. isaacii* PD618, *V. klebahnii* PD401, *V. tricorpus* PD593, and *V. zaregamsianum* PD739. Unless otherwise stated, fungi were maintained on potato dextrose agar (PDA) plates at room temperature in the dark.

The algal strains comprised three *Chlamydomonas reinhardtii* strains (SAG 11-32c, SAG 73.72, and CC 1690) and four additional *Chlamydomonas* species: *C. moewusii* (SAG 11-11), *C. incerta* (SAG 7.73), *C. applanata* (SAG 11-9), and *Edaphochlamys debaryana* (basionym *C. debaryana*, SAG 4.72) (22). Algal strains and species were purchased from the Culture Collection of Algae at the University of Göttingen (*Sammlung von Algenkulturen der Universität Göttingen*, SAG). Algal cultures were maintained in tris-phosphate medium with modified nitrogen sources (90% potassium nitrate, 10% ammonium) (TP10) (23) under a 14 h light/10 h dark photoperiod at a light intensity of ∼30 μmol photons m^−2^ s^−1^.

### Algal-fungal co-cultures

To establish co-cultures, fungal conidiospores were harvested from 5–7-day-old PDA plates, washed with deionized water, and resuspended in TP10 medium at 2 × 10^5^ spores mL^−1^. Algal suspensions were prepared from two-week-old pre-cultures and diluted in fresh TP10 medium to 2 × 10^4^ cells mL^−1^. Equal volumes of fungal and algal suspensions were combined, yielding final concentrations of 1 × 10^5^ spores mL^−1^ and 1 × 10^4^ cells mL^−1^, respectively, and distributed into 48-well plates. For co-cultures with different *Verticillium* and *Chlamydomonas* strains and species, algal final concentrations were 1 × 10^3^ cells mL^−1^, and suspensions were aliquoted into 96-well plates.

Axenic control cultures were prepared by mixing fungal or algal suspensions with TP10 medium at identical dilution ratios and distributing them as for co-culture treatments.

### Assessment of algal physiological responses

Physiological responses of algae to the presence of *Verticillium* were first assessed microscopically. Micrograph documentation of co-cultures was performed using a Motic AE2000 inverted microscope (Motic, Hong Kong) equipped with a MikroLive 6.4 MP CMOS camera (MikroLive, Oppenau, Germany). Autofluorescence of chlorophyll a/b was measured in 48-well plate cultures using a CLARIOstar Plus plate reader (BMG LABTECH, Offenburg, Germany) in 3 × 3 matrix mode with a 9 mm scan width and 2.4 mm focal height at excitation/emission wavelengths of 440–15/680–20 nm as a proxy for algal biomass. Photosynthetic activity was assessed by pulse-amplitude-modulated (PAM) fluorometry using an IMAGING-PAM system (IMAG-K7/IMAG-MAX/L, Heinz Walz GmbH, Effeltrich, Germany). Samples were dark-adapted for 15– 30 min prior to measurement. Measuring light, actinic light, and external light intensities were set to 6, 8, and 3, respectively. Saturating pulses were applied at an intensity of 7 with a duration of 240 ms. Photosynthetic efficiency (Fv/Fm) and chlorophyll fluorescence values were averaged across at least 24 technical replicates derived from three independent multi-well plates per condition.

### Fungal biomass quantification

For genomic DNA extraction, three multi-well plates per condition were frozen at −20 °C for at least 24 h and thawed at room temperature. To preserve sample integrity, plates were sealed with polyolefin cover foil (HJ-Bioanalytik, Erkelenz, Germany). Samples were then homogenized in each well using stainless steel beads (2.4 mm diameter; Omni International, Kennesaw, GA, USA) in 100 μL harvest buffer (10 mM Tris-HCl pH 8.0, 1 mM EDTA, 0.015% Tween 20) by vigorous shaking for 10 min at 12,000 rpm using a PSU-2T mini-shaker (SIA Biosan, Riga, Latvia). For each plate, homogenized material from all wells was pooled in 2 mL reaction tubes, collected by centrifugation, and snap-frozen in liquid nitrogen. Samples were further disrupted using a Retsch MM400 tissue lyser at maximum speed for ∼1 min and resuspended in 800 μL freshly prepared DNA extraction buffer. The extraction buffer consisted of basic extraction buffer (0.35 M sorbitol, 0.1 M Tris base, 5 mM EDTA, pH 7.5), nucleic lysis buffer (0.2 M Tris base, 50 mM EDTA, pH 7.5, 2 M NaCl, 2% CTAB), and 10% (w/v) sarkosyl in a 2.5:2.5:1 ratio. A spike-in plasmid (20 ng per sample; (24)) was added as an internal control. Samples were then incubated at 65 °C for at least 1 h, followed by addition of 0.5 volumes chloroform:isoamyl alcohol (24:1, v/v) and centrifugation at maximum speed for 15 min. The aqueous phase was transferred to a fresh tube, and DNA was precipitated by addition of one volume isopropanol followed by overnight incubation at −20 °C. DNA was pelleted by centrifugation at 10,000 rpm for 10 min, washed with 70% ethanol, air-dried for 5–10 min, and resuspended in Milli-Q water. Quantification of fungal biomass was performed by real-time qPCR using SsoAdvanced Universal SYBR Green Supermix (Bio-Rad, Hercules, CA, USA) with primers targeting *VdITS1* (F 5′-AAAGTTTTAATGGTTCGCTAAGA-3′, R 5′-CTTGGTCATTTAGAGGAAGTAA-3′) and the spike-in plasmid (F 5′-TTTCTTTTCCAAGGTTTGTGC-3′, R 5′-AACATTTACCCTGCTTGTAGCTCT-3′). PCR cycling conditions were: 98 °C for 3 min, followed by 36 cycles of 98 °C for 10 s and 60 °C for 15 s. A melt curve analysis (65–95 °C, 0.5 °C increments) was performed to verify amplification specificity. *V. dahliae ITS1* levels were normalized to the spike-in plasmid using the ΔCt method.

### Chemotaxis assay

Chemotaxis was assessed by quantifying *C. reinhardtii* cells that migrated into glass capillaries containing potential attractants (19). Capillaries (3 μL; Drummond Microcaps®, Drummond Scientific, Broomall, PA, USA) were filled with fresh TP10 medium or algal culture filtrate (controls), or with supernatant from a 7-day-old *V. dahliae* culture grown in TP10 medium. Loaded capillaries were placed upright in 2 mL reaction tubes containing 100 μL of a dense *C. reinhardtii* suspension, such that cells had to swim against gravity to enter the capillaries. Samples were incubated in the dark at 25 °C for 5 h. Capillary contents were then transferred to 27 μL PBS supplemented with 3 μL Lugol’s solution (Sigma-Aldrich, Darmstadt, Germany), and migrated cells were quantified using a Neubauer counting chamber.

### Stress assays

Algal–fungal co-cultures and monocultures were established as described above. After 3 days of growth, cultures were exposed to osmotic stress by addition of 300 mM mannitol or 100 mM NaCl. To assess responses to biotic stress, cultures were treated with 35 μM of the cyclic lipopeptide orfamide A (Santa Cruz Biotechnology, TX, USA), a toxin produced by the algicidal bacterium *Pseudomonas protegens* Pf-5 (25). Effects on algal physiology were assessed after 4 days of treatment by measuring chlorophyll fluorescence as a proxy for algal biomass, as described above.

### Statistics

Statistical analyses were performed in R (version 4.5.2) using *lmerTest* within the *lme4* package. Unless otherwise indicated, linear mixed-effects models were fitted with ‘condition’ (axenic or co-culture) as a fixed effect and ‘experimental replicate’ as a random effect to account for the nested structure of the data (multiple wells measured for each condition within each plate). Plates were treated as experimental units. P-values were obtained using Satterthwaite’s approximation as implemented in the *lmerTest* package. Results were consistent with analyses performed on plate-level means (n = 3 independent plates per condition).

## RESULTS

### A mutualistic interaction between *C. reinhardtii* and *V. dahliae*

*C. reinhardtii* and *V. dahliae* both occur in terrestrial habitats where they interact with diverse soil microbiota (20,23,26,27). To explore potential interactions between these organisms, we established a microtiter plate co-culture assay under defined laboratory conditions in which *C. reinhardtii* strain CC1690 and *V. dahliae* strain JR2 can be cultivated together in a carbon-free growth medium that only supports limited fungal growth (Figure 1A, B). After 5 days of culturing, both organisms had grown in axenic cultures and algal cells clearly flocculated around fungal hyphae in co-cultures, resulting in close physical contact (Figure 1B). Initially, cultures were kept at relatively low light intensities (∼40 μmol photons m^-2^ s^-1^) (28). Under these conditions, chlorophyll fluorescence intensity was significantly higher in co-cultures compared to axenic algal controls (Figure 1C), suggesting that the presence of *V. dahliae* positively affects algal physiology. However, under natural conditions, both organisms are likely exposed to much lower light intensities. Therefore, we subsequently tested whether the fungus similarly affects *C. reinhardtii* under very low light conditions (∼20 μmol photons m^-2^ s^-1^). Chlorophyll fluorescence measurements showed that fluorescence intensity was significantly enhanced in the presence of *V. dahliae* compared to axenic control cultures (Figure 1D), indicating that *C. reinhardtii* benefits from fungal presence also in very low light. Since bulk chlorophyll fluorescence can be influenced by changes in chlorophyll content or photosynthetic physiology, photosynthetic activity was measured by pulse-amplitude-modulation (PAM) fluorometry. In contrast to chlorophyll fluorescence, photosynthetic efficiency was similar in co-cultures and in algal monocultures in very low light (Figure 1E), suggesting that the elevated chlorophyll fluorescence intensity measured in co-cultures primarily reflects increased algal biomass rather than enhanced photosystem II performance per cell. Consistent with this interpretation, fluorescence measurements across algal cultures inoculated with different initial cell densities showed dose-dependent fluorescence intensities that increased over time (Supplementary Figure S1), supporting the use of chlorophyll fluorescence as a proxy for algal biomass under these assay conditions. Concomitantly, fungal biomass was significantly enhanced in co-culture with *C. reinhardtii* compared to fungal monocultures (Figure 1F). Together, these results indicate that the interaction between *C. reinhardtii* and *V. dahliae* is mutually beneficial.

**Figure 1.**
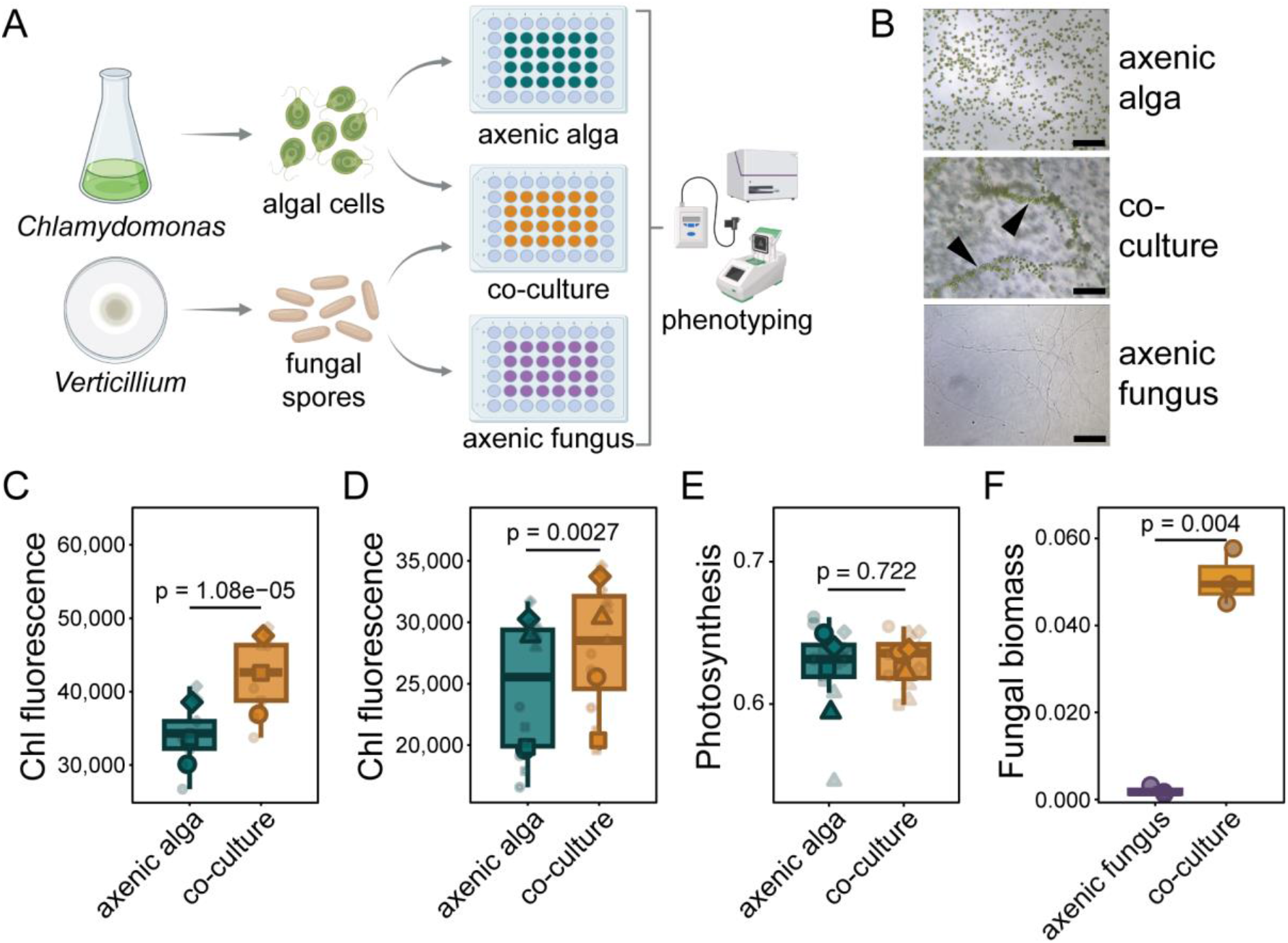
Enhanced growth in *C. reinhardtii–V. dahliae* co-cultures. (A) Schematic representation of the experimental setup. *C. reinhardtii* cells were co-inoculated with *V. dahliae* spores at a 1:10 ratio in carbon-free growth medium in 48-well plates and grown under 14h/10h light/dark conditions at 20°C. Axenic *C. reinhardtii* and *V. dahliae* cultures were prepared as controls. Figure generated with BioRender.com. (B) Following 5 days of co-culturing, *C. reinhardtii* cells clearly flocculated around *V. dahliae* hyphae (black arrowheads) visible by light microscopy. Size bar = 100 µm. After 7 days of culturing under low light conditions at ∼40 μmol photons m^-2^ s^-1^ (C) or very low light conditions at ∼20 μmol photons m^-2^ s^-1^ (D), chlorophyll fluorescence was significantly enhanced in presence of the fungus. (E) In contrast, measurements of maximum quantum efficiency of photosystem II (F_v_/F_m_) as indicator of photosynthetic activity showed no difference between axenic *C. reinhardtii* cultures or co-cultures with *V. dahliae*. (C-E) Small symbols represent independent plates, and large symbols indicate means (n = 3 plates per condition). Different symbol shapes represent independent experiments. Statistical significance was assessed using linear mixed-effects models. (F) Quantification of fungal biomass by quantitative real-time PCR on genomic DNA showed that the growth benefit is mutual with the fungus displaying enhanced biomass in presence of the alga compared to axenic control cultures. Species-specific primers targeting *V. dahliae ITS* were used for quantification with primers against a spike plasmid used for calibration. Graph shows data of one of three independently performed experiments with similar results. Statistically significant difference between treatments was determined by an unpaired, two-sided Student’s t-test with n = 3 per condition.

### Recruitment to fungal hyphae and direct contact enable the algal growth benefit

Algal–fungal interactions are generally characterized by direct physical contact between the partners (18,29). Consistent with this, microscopic examination of liquid co-cultures revealed clear physical association between *C. reinhardtii* and *V. dahliae*, with algal cells in close contact with fungal mycelium (Figure 1B, Figure 2A).

**Figure 2.**
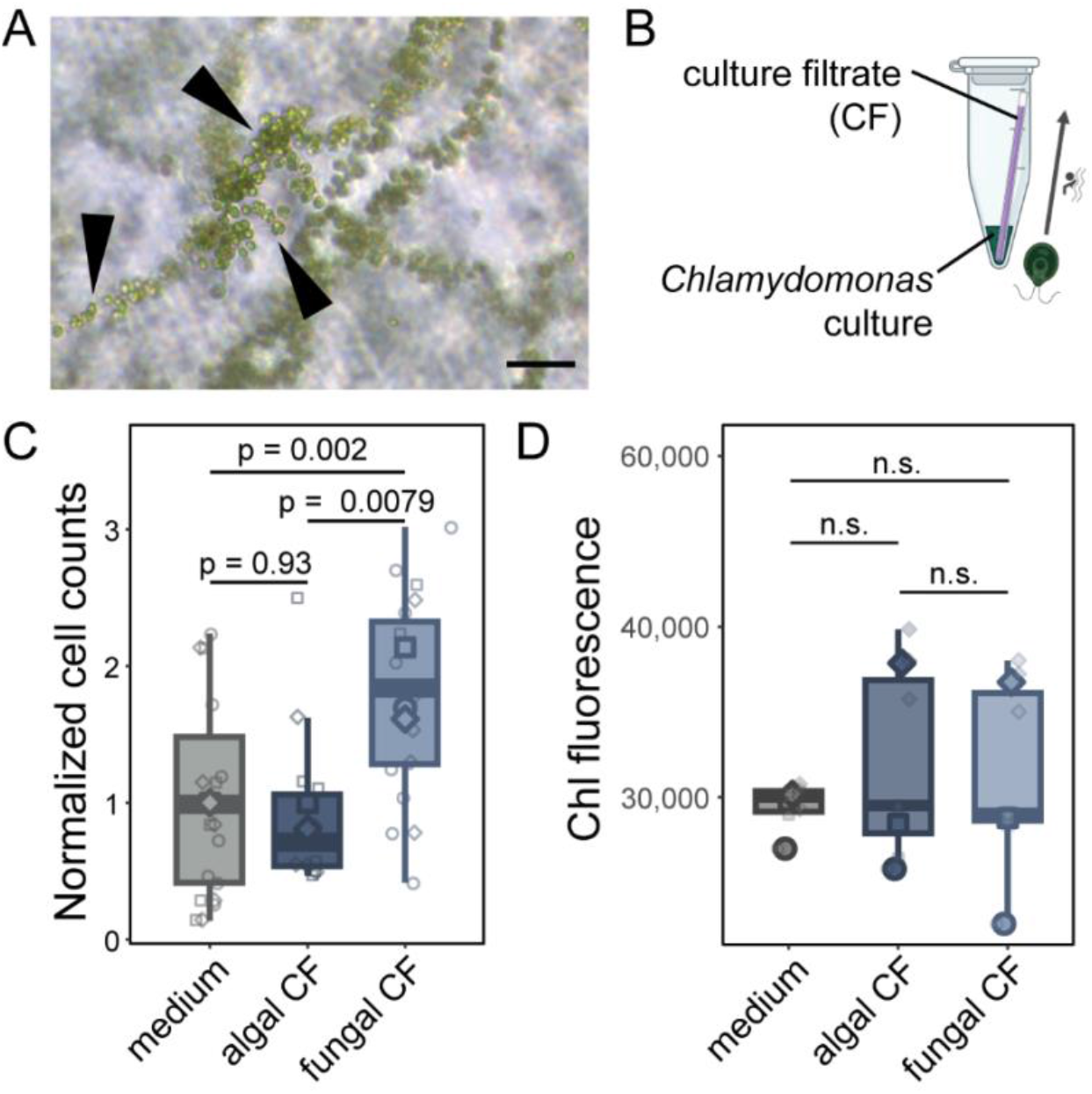
Chemotactic recruitment of *C. reinhardtii* to fungal hyphae and requirement for physical contact in the mutualistic interaction. (A) Micrograph showing *C. reinhardtii* cells in direct physical contact (black arrowheads) with *V. dahliae*. Size bar = 50 µm. (B) *V. dahliae* attracts *C. reinhardtii* cells. Capillaries filled with fresh culture medium (TP10), algal culture filtrate (CF), or fungal CF were placed into tubes containing *C. reinhardtii* suspensions. (C) Following 5 h of incubation, numbers of *C. reinhardtii* cells that migrated into capillaries containing fungal CF were significantly higher than for the algal CF and medium controls. (D) Fungal exudates do not enhance *C. reinhardtii* growth. *C. reinhardtii* cells were cultured in fresh culture medium (TP10), supplemented with algal CF or fungal culture filtrate CF. Algal growth was determined by chlorophyll fluorescence measurement after 7 days of culturing. Small symbols represent independent plates, and large symbols indicate means (n = 3 plates per condition). Different symbol shapes represent independent experiments. Statistical significance was assessed using linear mixed-effects models.

To determine whether algal cells are actively recruited to the fungus, we tested whether *C. reinhardtii* exhibits chemotactic responses to fungal exudates. In a chemotaxis assay, glass capillaries were filled with fungal culture filtrate and placed into tubes containing algal suspensions (Figure 2B). Capillaries filled with algal culture filtrate or fresh culture medium were used as negative controls. After 5 hours, significantly more *C. reinhardtii* cells had swum into capillaries containing fungal culture filtrate compared with those containing algal culture filtrate or fresh medium (Figure 2C), indicating that diffusible fungal exudates attract algal cells.

We next assessed whether such diffusible signals are sufficient to induce the algal growth benefit observed in co-culture. Culture filtrates were collected from axenic fungal and algal cultures and added to fresh culture medium. *C. reinhardtii* cells were then grown either in fresh medium alone or in medium supplemented with the respective culture filtrates. Chlorophyll fluorescence measurements showed that algal biomass did not differ between the cultures grown in fresh medium and cultures supplemented with *C. reinhardtii* culture filtrate (Figure 2D). Interestingly, in contrast to direct co-cultures with the fungus, algal biomass in cultures treated with fungal culture filtrate did not differ from the controls (Figure 2D). These results indicate that fungal exudates alone are insufficient to reproduce the algal growth benefit observed in co-culture. Together, these findings show that diffusible fungal signals recruit *C. reinhardtii* cells toward fungal hyphae, while direct physical contact with the fungus appears necessary for the enhanced algal growth observed in co-culture.

### Fungal association supports algal performance under stress

Terrestrial microalgae frequently experience fluctuations in water availability that impose osmotic stress. In lichen symbioses, the fungal partner contributes to water retention and buffering of environmental conditions, which can reduce osmotic stress experienced by the photobiont (5). This is in accordance with the general notion that mutualistic microbial interactions can enhance stress tolerance of the interacting partners (30). To determine whether the interaction with *V. dahliae* similarly improves stress resilience of *C. reinhardtii*, the alga was grown axenically or in co-culture with the fungus. After three days of growth, cultures were exposed to abiotic stress induced either by NaCl, which imposes combined ionic and osmotic stress, or by mannitol, which induces non-ionic osmotic stress. Both stressors were applied at concentrations previously shown to substantially reduce algal growth without causing cell death (31). Following four days of stress exposure, algal growth was assessed by measuring chlorophyll fluorescence. Under both stress conditions, the biomass of *C. reinhardtii* remained significantly higher in co-culture with *V. dahliae* compared with axenic controls (Figure 3A,B). These results indicate that the growth benefit conferred by the fungal partner persists under abiotic stress conditions.

**Figure 3.**
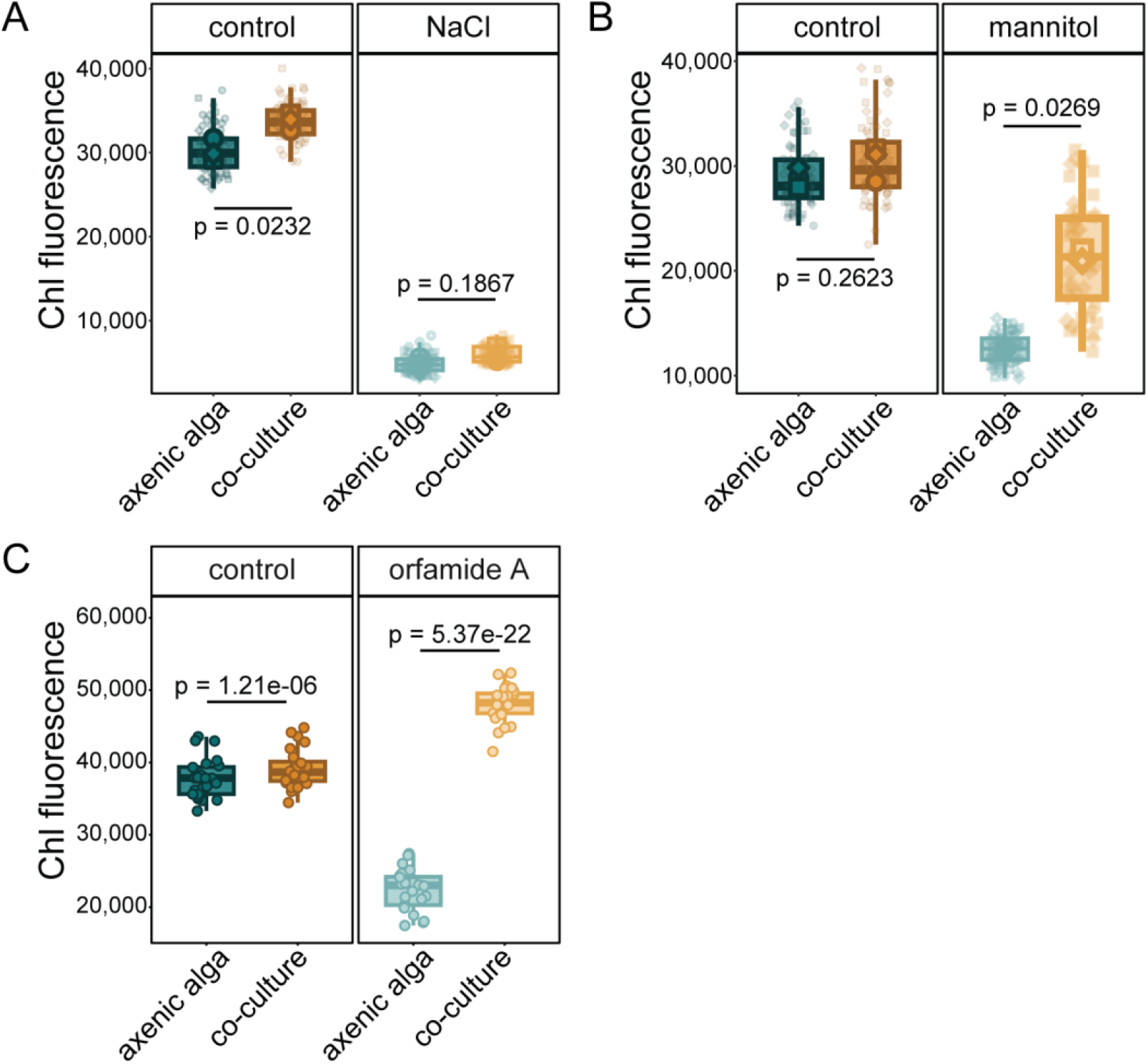
Algal growth benefit is maintained under diverse stress conditions. To test whether the mutualistic interaction between *C. reinhardtii* and *V. dahliae* affects algal stress resilience, 3-day-old axenic algal cultures or co-cultures with the fungus were treated with 100 mM NaCl (A), 300 mM mannitol (B) or 35 µM orfamide A (C). After 4 days of treatment, the effect on algal biomass was assessed through chlorophyll fluorescence measurements, showing that the presence of the fungus mitigates the effect of both abiotic stress and microbial antagonism on the alga. Statistically significant differences between treatments according to an unpaired two-sided Student’s t test on n=3 plates per condition. Data are shown from one of three independent experiments with similar results.

To quantify the extent to which fungal association buffered stress effects, we calculated the percent change of algal chlorophyll fluorescence under stress compared with unstressed controls in absence and presence of *V. dahliae*. While the magnitude of the NaCl-induced biomass reduction was comparable in both culture conditions (Supplementary Figure S2A), the mannitol-induced reduction of algal biomass was considerably smaller in co-culture than in axenic cultures (Supplementary Figure S2B), supporting the conclusion that *V. dahliae* selectively enhances abiotic stress tolerance in *C. reinhardtii*.

In addition to abiotic stress, algae are frequently exposed to antagonistic microorganisms that release algicidal compounds in natural environments. One such compound is the cyclic lipopeptide orfamide A, produced by the bacterium *Pseudomonas protegens* Pf-5, which contributes to the detrimental effect of this bacterium on *C. reinhardtii* (25). To determine whether the fungal partner can also mitigate the effects of such microbial toxins, axenic algal cultures and algal–fungal co-cultures were exposed to orfamide A. Treatment with orfamide A strongly reduced the biomass of axenic *C. reinhardtii* cultures relative to untreated controls (Figure 3C). In contrast, algal cultures grown in association with *V. dahliae* displayed significantly higher chlorophyll fluorescence levels under toxin exposure even surpassing fluorescence intensity values measured in unstressed axenic control cultures (Figure 3C). These results indicate that the beneficial interaction with the fungus mitigates the detrimental effect of the algicidal compound and supports algal performance during microbial antagonism.

### Interaction outcomes depend on algal–fungal pairings

To test whether the capacity to confer a growth benefit to *C. reinhardtii* is a conserved trait within the *Verticillium* genus, we first co-cultured our focal *C. reinhardtii* strain CC1690 with different *V. dahliae* strains that display varying degrees of virulence on vascular plant hosts. In addition to *V. dahliae* JR2, co-cultures with the strains DVD3, TO22 and CQ2 resulted in a highly significant increase in chlorophyll fluorescence intensity (Figure 4A) corresponding to a 0.2-0.4 log_2_ fold-change compared to axenic controls (Figure 4C), indicating that *C. reinhardtii* biomass is also enhanced in the presence of these strains. The strains VdLs17, 85S, and V700 caused a moderate increase in biomass and only strain Vd39 had a significant negative effect on *C. reinhardtii* biomass (Figure 4A,C).

**Figure 4.**
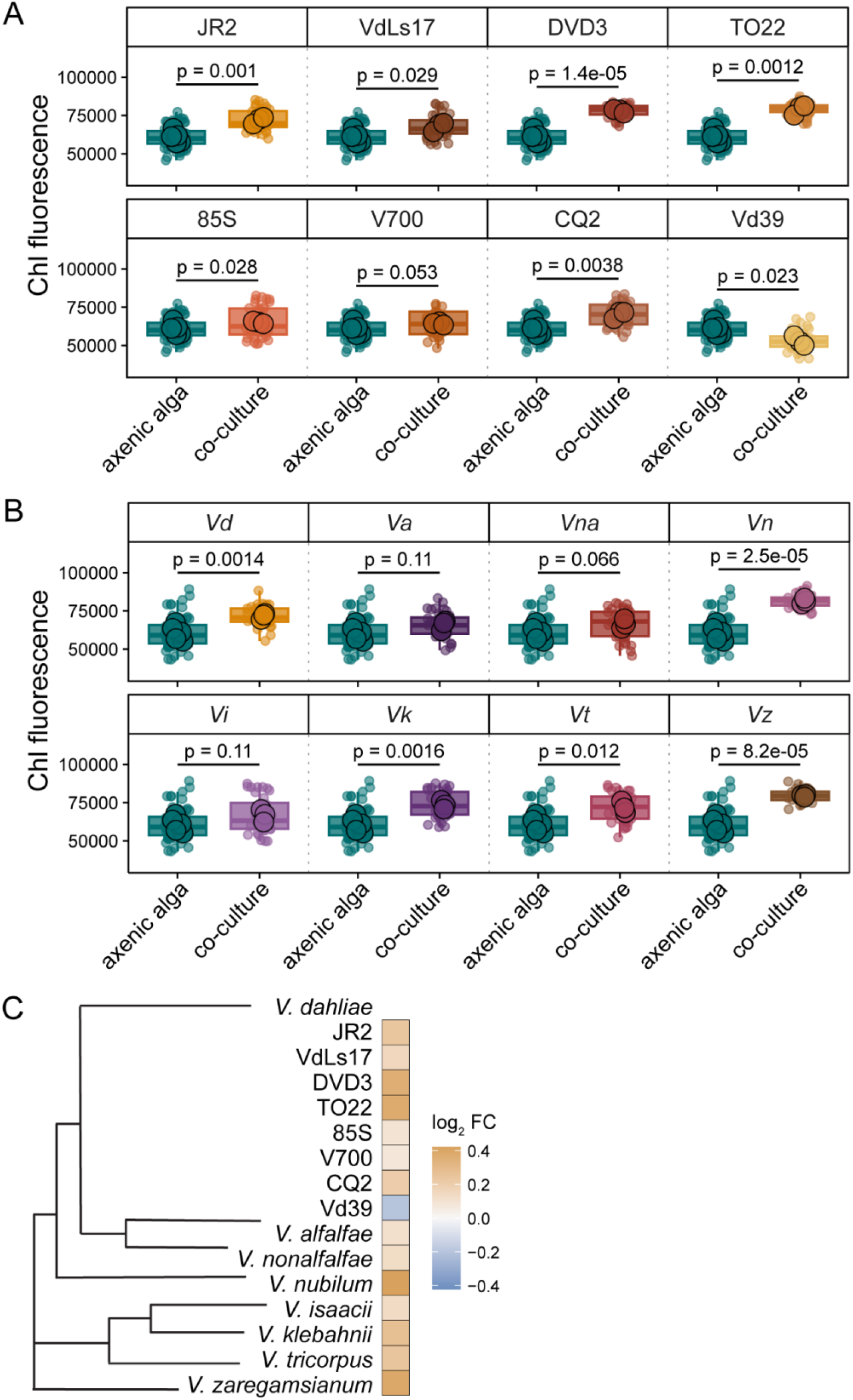
Positive effects on algal growth are broadly conserved across the *Verticillium* genus. Chlorophyll fluorescence of *C. reinhardtii* strain CC1690 grown axenically or in co-culture with indicated *V. dahliae* strains (A), or in co-culture with different *Verticillium* species (B) for 11 days. In (A) and (B), individual points represent measurements from single wells and large circles represent plate means. Statistical significance was assessed using two-sided t-tests on plate means, treating individual plates as independent experimental units (n = 3 per condition). Axenic algal controls were included in each experiment and are shown in each facet for comparison. The *V. dahliae* entry in (B) corresponds to strain JR2 in (A). (C) Phylogenetic relationships among *Verticillium* strains and species used in this study, with corresponding effects on algal chlorophyll fluorescence shown as a heatmap. Colours represent the log_2_ fold change in fluorescence in co-culture relative to axenic conditions, with positive values indicating increased fluorescence and negative values indicating decreased fluorescence. Data are shown from one of three independent experiments with similar results. *Vd*: *V. dahliae, Va*: *V. alfalfae, Vna*: *V. nonalfalfae, Vn*: *V. nubilum, Vi*: *V. isaacii, Vk*: *V. klebahnii, Vt*: *V. tricorpus, Vz*: *V. zaregamsianum*.

We then co-cultured *C. reinhardtii* CC1690 with different *Verticillium* species whose lifestyles range from (opportunistic) plant parasitism to saprotrophism. Interestingly, all *Verticillium* species caused moderate to strong increases of fluorescence intensity in co-cultures compared to algal monocultures (Figure 4B). The strongest effects were exerted by *V. nubilum* and *V. zaregamsianum* with log_2_ fold-changes of 0.42 and 0.39, respectively (Figure 4C). These findings indicate that the beneficial effect on *C. reinhardtii* CC1690 growth extends beyond *V. dahliae* and is broadly conserved across all *Verticillium* species, irrespective of their lifestyle or phylogenetic position.

We then asked whether different *Chlamydomonas reinhardtii* strains are similarly positively affected by *V. dahliae*. To test this, we monitored the growth of the two *C. reinhardtii* strains SAG 11-32c and SAG 73.72 in addition to CC1690 in co-culture with our focal *V. dahliae* strain JR2. Apart from CC1690, only SAG 73.72 showed enhanced chlorophyll fluorescence in presence of the fungus compared to axenic algal cultures, whereas SAG 11-32c was not affected at all (Figure 5A,C), suggesting that *C. reinhardtii* strains are differentially affected by the fungus.

**Figure 5.**
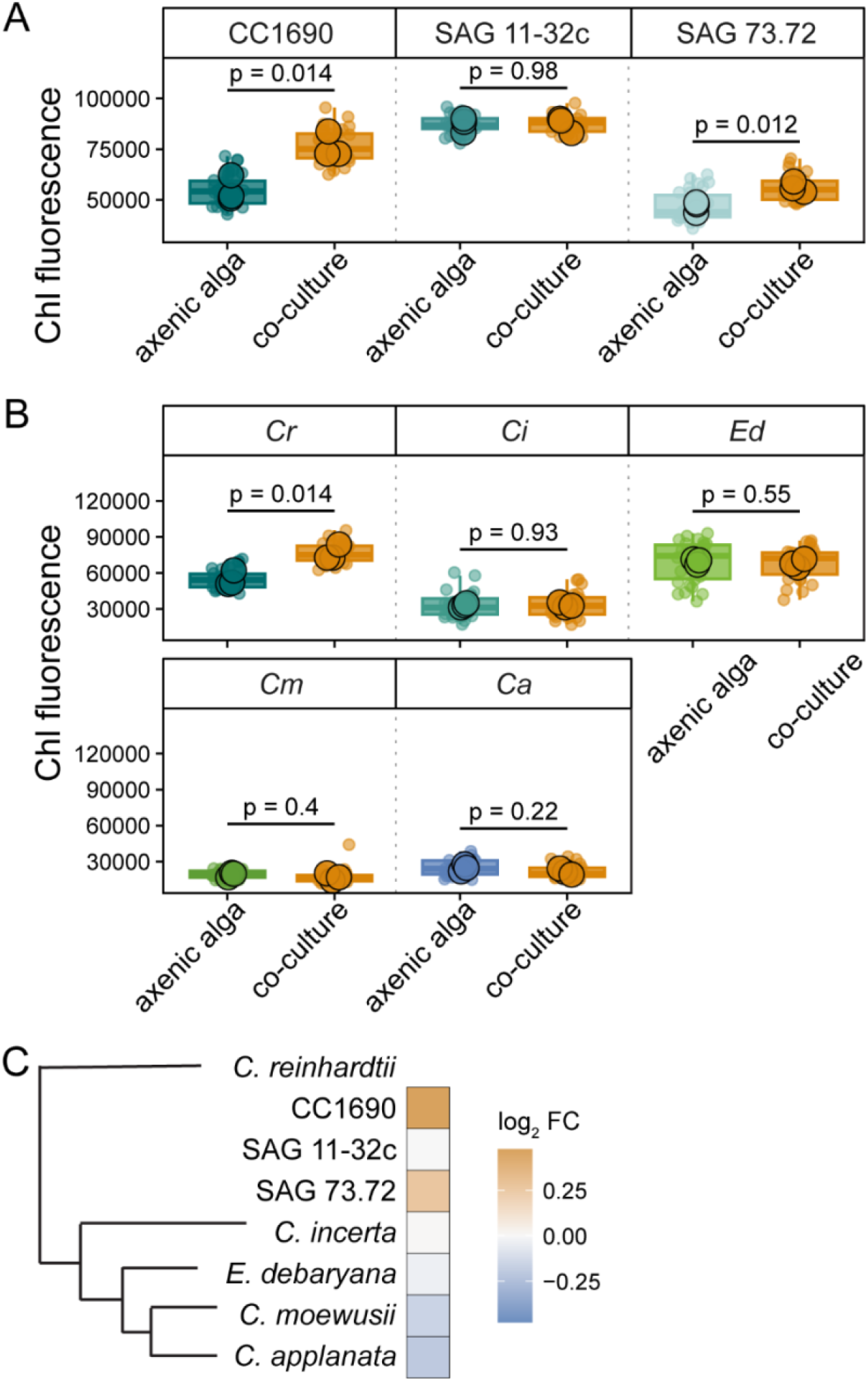
Algal growth responses to *V. dahliae* vary across *Chlamydomonas* lineages. Chlorophyll fluorescence of *C. reinhardtii* strains CC1690, SAG 11-32c, SAG 73.72 (A) and of different *Chlamydomonas* species (B) grown axenically or in co-culture with *V. dahliae* JR2 for 11 days. In (A) and (B), individual points represent measurements from individual wells and large circles indicate plate means. Statistical significance was assessed using two-sided Student’s t-tests on plate means, treating individual plates as independent experimental units (n = 3 per condition). The *C. reinhardtii* entry in (B) corresponds to strain CC1690 in (A). (C) Phylogenetic relationships among *Chlamydomonas* strains and species included in this study, with corresponding effects of fungal co-culture on chlorophyll fluorescence shown as a heatmap. Colors represent the log_2_ fold change in fluorescence in co-culture relative to axenic conditions. Data are shown from one experiment representative of three independent experiments. *Cr*: *C. reinhardtii, Ci*: *C. incerta, Ed*: *E. debaryana, Cm*: *C. moewusii, Ca*: *C. applanata*.

Notably, when we co-cultured other *Chlamydomonas* species with *V. dahliae* JR2, we did not observe an increase in chlorophyll fluorescence for any of the partner combinations (Figure 5B,C), indicating that the ability to engage in mutualism with this fungus may be restricted to *C. reinhardtii*. Moreover, the effect of *V. dahliae* on *Chlamydomonas* spp. became increasingly negative as indicated by decreasing log_2_ fold-changes correlating with their phylogenetic distance to *C. reinhardtii* (Figure 5C). These findings indicate that algal growth responses to *V. dahliae* vary between *Chlamydomonas* species and that species-specific traits influence the interaction outcome.

## DISCUSSION

Interactions between algae and fungi are widespread across diverse ecosystems, yet the molecular mechanisms that govern these associations remain poorly understood. Progress in this area has been limited, in part, by the complexity of natural systems and the lack of experimentally tractable models. Here, we establish a controlled synthetic system between the model species *C. reinhardtii* and *V. dahliae*. Surprisingly, this interaction resulted in growth benefits for both partners reminiscent of mutualistic interactions. *V. dahliae* is best known as a pathogen infecting hundreds of plant species (26). In plants, pathogenicity is associated with vascular colonization, secretion of cell wall-degrading enzymes, and production of phytotoxic compounds (32). However, co-cultivation of *C. reinhardtii* with *V. dahliae* did not result in detectable alterations in algal cell morphology but instead led to enhanced algal proliferation. These findings suggest that fungal pathogenic mechanisms are not activated or are not effective in this interaction context. Importantly, our observations are consistent with earlier studies showing that *V. dahliae* is not restricted to a pathogenic lifestyle and can occur as an endophyte in several plant hosts (33–36), indicating that interaction outcomes with plants and algae are context-dependent. These observations raise the question of how this interaction is established and maintained at the cellular level.

Physical contact between partners is a defining feature of many algal–fungal symbioses. In lichens and related associations, close spatial organization of fungal and algal cells is thought to facilitate coordinated interactions between partners (7,29,37). Direct cell–cell contact has also been reported in synthetic algal–fungal systems (15,18). However, not all systems require physical proximity: algal growth can be enhanced in the absence of direct contact when diffusible cues are exchanged between *C. reinhardtii* and *A. nidulans* (19), and long-term associations with *A. infectora* occur without direct contact despite embedding in an extracellular matrix (17,37). In contrast, our results indicate that diffusible *V. dahliae* factors alone are insufficient to promote *C. reinhardtii* growth, as exposure to fungal culture filtrates did not recapitulate the growth benefit observed in co-culture. Instead, our findings point to a requirement for close association between the partners, even though we did not observe the formation of morphologically complex structures typical of lichens (37). Consistent with this, *C. reinhardtii* actively responds to fungal exudates and exhibits directed movement toward fungal hyphae, similar to its behaviour toward *A. nidulans* (19). Together, these findings suggest that while diffusible signals may mediate partner attraction, close physical association is likely required for the establishment of a functional mutualistic interaction in our system. A key question is whether such close associations translate into functional advantages for the interacting partners.

Fungal mutualistic interactions often confer enhanced stress resilience to their partners (5,38). In line with this, we observed that the association with *V. dahliae* enhances the resilience of *C. reinhardtii* under abiotic stress conditions, indicating that the interaction confers functional benefits. Such effects may arise through microenvironmental buffering or modulation of stress-responsive signalling pathways between partners (30,39–41). Protective effects of microbial interactions have also been described in the context of biotic stress. In synthetic consortia, *C. reinhardtii* can be shielded from algicidal compounds by associated microbes. For instance, in association with *A. nidulans*, algal cells are protected from the toxin azalomycin F produced by *Streptomyces iranensis* through detoxification by fungal lipid compounds (19), while the bacterium *Mycetocola lacteus* protects *C. reinhardtii* from the cyclic lipopeptide orfamide A produced by *P. protegens* via enzymatic inactivation (42). Consistent with these observations, we find that *V. dahliae* mitigates the toxic effects of orfamide A on *C. reinhardtii*. Whether this protection involves direct detoxification, chemical modification of the toxin, or indirect effects such as altered algal physiology remains to be determined. Together, these findings suggest that the *Chlamydomonas–Verticillium* interaction enhances algal fitness under both abiotic and biotic stress conditions, highlighting its potential ecological relevance in fluctuating environments. However, whether such beneficial interactions are consistently established or vary depending on the identity of the interaction partners remains unclear.

Our comparative analysis across multiple *Chlamydomonas* and *Verticillium* strains reveals that interaction outcomes are strongly influenced by partner identity but are not symmetric across the two partners. While most *Verticillium* strains and species conferred a growth benefit to *C. reinhardtii*, this effect was not consistently observed across different *Chlamydomonas* strains and species, indicating that the capacity to promote algal growth is broadly conserved within the *Verticillium* genus but restricted on the algal side. These findings suggest that interaction outcomes are primarily shaped by species-specific properties of the algal partner rather than reflecting a general incompatibility between algae and fungi. In natural communities, microbial interactions are highly context-dependent, with outcomes ranging from cooperation to antagonism depending on environmental conditions and the physiological and genetic compatibility of the partners involved (43–46). In algal–fungal systems, such variation may arise from differences in the ability to perceive and respond to partner-derived signals, sensitivity to secreted compounds, or the establishment of compatible interaction interfaces. The observed patterns therefore suggest that partner recognition and compatibility play a central role in determining whether interactions result in mutualistic or neutral outcomes. In this context, the *C. reinhardtii–V. dahliae* system provides a tractable platform to dissect the molecular mechanisms underlying partner specificity in algal–fungal associations.

Reciprocal exchange of nutrients or metabolites is a common feature of mutualistic interactions, including algal–fungal associations such as lichens (5). Experimental systems have shown that such exchanges can support mutualistic growth under laboratory conditions (15,17). In our system, the mutual growth enhancement observed for both partners is consistent with the possibility of reciprocal resource exchange, for example through the provision of photosynthetically derived carbon to the fungus and increased availability of dissolved CO_2_ to the alga. However, the mechanisms underlying this interaction remain to be determined, and future work will be required to establish whether and how metabolites are exchanged between partners. Together, these findings highlight both the complexity of algal–fungal interactions and the need for experimentally tractable systems to dissect their underlying mechanisms.

Together, our findings establish the *Chlamydomonas reinhardtii–Verticillium dahliae* system as a tractable model for studying algal–fungal interactions under controlled conditions. By combining genetic accessibility with a reproducible and stable mutualistic interaction phenotype, this system enables mechanistic investigation of processes such as metabolite exchange, partner recognition, and signalling, as well as the determinants of interaction outcomes. Future work may reveal the molecular basis of algal–fungal communication and how environmental context shapes interaction dynamics, advancing our understanding of these interactions across ecological and evolutionary scales.

## ACKNOWLEDGEMENTS

We thank Gerlind Bauerecker for technical assistance and Bart P. H. J. Thomma for critically reading the manuscript.

## AUTHOR CONTRIBUTIONS

H.R. and Z.P. conceived the study. Z.P. and U.N. performed experiments. Z.P., U.N., and H.R. developed methodology and analysed data. Z.P. and H.R. wrote the manuscript with input from all authors. H.R. supervised the study and acquired funding.

