## Supplemental Material for "The model alga *Chlamydomonas reinhardtii* forms mutualistic interactions with *Verticillium* fungi"

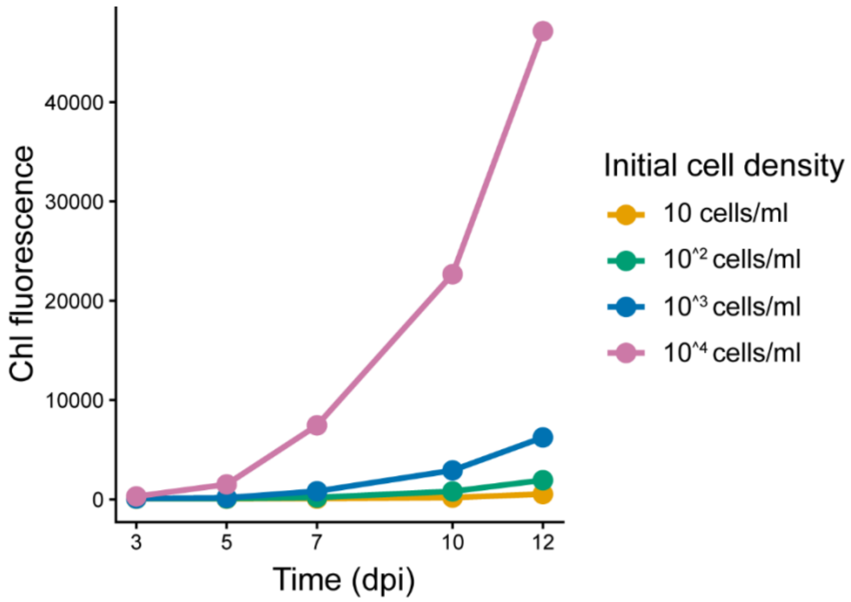


**Supplementary Figure S1. Chlorophyll fluorescence correlates with algal cell density.** Axenic *C. reinhardtii* cultures were inoculated at increasing initial cell densities and monitored over time. Higher starting densities resulted in proportionally higher fluorescence signals, supporting the use of chlorophyll fluorescence as a proxy for algal biomass.

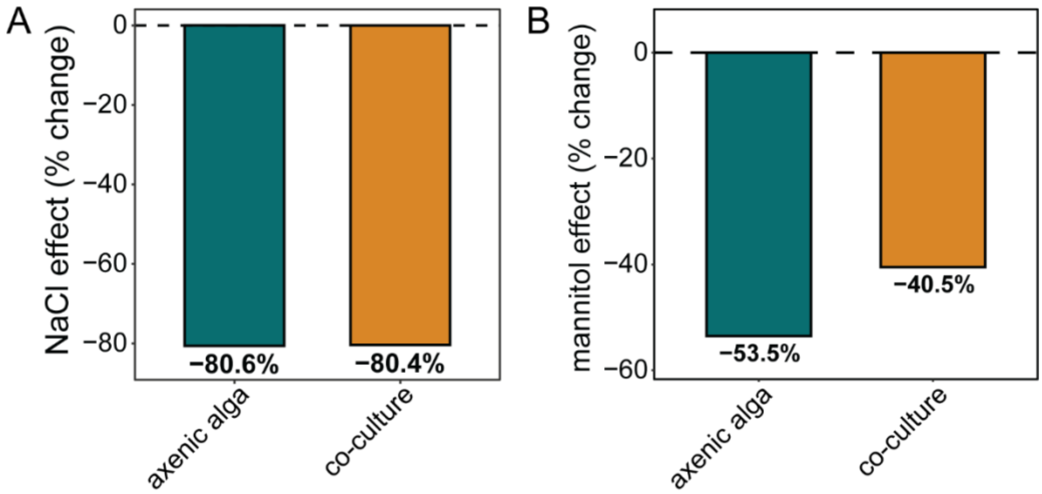


**Supplementary Figure S2. Effect sizes of osmotic stress treatments in axenic and co-culture conditions.** Percent change in chlorophyll fluorescence under NaCl (A) and mannitol (B) treatment relative to untreated controls. Values indicate the magnitude of stress-induced fluorescence intensity reduction in axenic *C. reinhardtii* cultures and in co-culture with *V. dahliae* from 3 independently performed experiments.
